# Directional bone matrix mineralization in the CAM assay is governed by vascular integration and matrix remodeling

**DOI:** 10.64898/2026.05.29.728599

**Authors:** Shola Onissema Karimu, Stephan M. Sutter, Siyoung Choi, Shuofei Sun, Jonathan Butcher, Lara A. Estroff, Claudia Fischbach

## Abstract

Bone matrix mineralization plays an important role in maintaining bone health but is a highly dynamic process for which few physiologically relevant model systems exist. The chorioallantoic membrane (CAM) assay is one such assay and has been used to study biomineralization; however, the mechanisms controlling mineral deposition in the CAM assay remain poorly understood. Here, we implanted decellularized, organic bone matrices onto the CAM and investigated their mineralization using a combination of micro-computed tomography, Raman spectroscopy, scanning electron microscopy, and histochemistry. These studies revealed increased mineralization and microvessel density on the shell-facing interface of the implanted matrices, while their embryo-facing counterpart lacked mineralization but was densely infiltrated by cells. Seeding organic bone matrices with mesenchymal stromal cells (MSCs) prior to CAM-implantation prevented mineralization by altering matrix micro- and nanostructure and limiting vessel integration. Collectively, our results suggest that mineral precursors from the eggshell and vasculature combine to mineralize organic bone matrix in the CAM assay, while proteolytic remodeling by invaded or implanted stromal cells inhibits that process. Our findings provide critical new insights into the interplay between acellular and cellular drivers of bone matrix mineralization, and will inform future studies of biomineralization using the CAM assay.

## Introduction

Bone matrix mineralization is critical for the skeleton’s unique mechanical properties, maintenance of systemic calcium and phosphate homeostasis, and regulation of cell signaling. The loss of bone mineral density or quality, which is common during aging, following fracture healing, or in diseases such as osteoporosis and cancer, is accordingly associated with poor patient prognosis and significant socioeconomic consequences^1–5^. Current therapeutic strategies to compensate for properties otherwise mediated by bone matrix mineralization include drug treatment or implantation of biomaterials or bone grafts. While these approaches can alleviate symptoms (*e.g.*, pain) and restore some of bone’s intrinsic functionalities (*e.g.*, mechanical stability), they are accompanied by negative side effects and unsatisfactory long-term prognosis^6,7^. Better understanding of the mechanisms that promote or inhibit mineralization of physiological bone matrix will help identify novel therapeutic targets that can address these limitations.

Bone matrix is largely composed of densely packed collagen fibrils reinforced with hydroxyapatite (HA; Ca_10_(PO_4_)_6_(OH)_2_) nanoparticles, which together endow bone with its unique mechanical properties^8,9^. The mineral content of bone matrix is regulated by the balance between bone formation and resorption – collectively known as remodeling. Importantly, shifts in physiological calcium and phosphate concentrations influence bone health and modulate cellular activity^10^. Research into conditions associated with reduced bone mineral density has revealed important insights into how overactive osteoclasts and/or impaired mesenchymal stem cells (MSCs) compromise bone matrix mineralization^11,12^. However, the role that bone matrix plays in this process remains poorly understood – a crucial missing piece, given that bone matrix mineralization is known to promote MSC differentiation into osteoblasts^8,13^ which, in turn, deposit the collagen type I-rich osteoid that provides the template for subsequent mineralization^8^. In addition, we recently showed that lack of bone matrix mineralization promotes a profibrotic rather than osteogenic MSC phenotype with possible functional consequences on bone matrix mineralization^14^. However, which potential effects the interplay between bone matrix and MSCs have on mineralization is unclear. Studying these connections requires model systems and analytical approaches that allow investigating these connections in a physiologically relevant environment and ensure the formation of chemically defined bone mineral.

Investigating the combined effects of bone matrix mineral content and MSCs *in vivo* is challenging, as either of these components is difficult to selectively control with animal studies. *In vitro* platforms including scaffold-based cultures and hydrogel systems have emerged as an alternative method^6,15–20^ but often lack key physiological conditions that influence mineralization including (*i*) relevant ion concentrations, (*ii*) the 3D nano- and micro-architecture of organic bone matrix necessary for bone-like intrafibrillar mineralization, and (*iii*) mass transport conditions that mimic the supply of small molecules as well as growth factors and cytokines involved in bone formation. Thus, alternative model systems are needed to explicitly study bone matrix mineralization under well-defined and yet physiologically relevant conditions.

The chorioallantoic membrane (CAM) assay presents a promising tool to examine bone matrix mineralization under *in vivo*-like conditions. The CAM contains a rich vascular network that connects with the embryonic circulatory system. It facilitates gas exchange^21^ between the embryo and the external environment and simultaneously enables skeletal development of the developing chick by transporting calcium ions mobilized from the eggshell^22^. Moreover, the CAM vasculature provides direct access to phosphate ions, largely derived from the yolk phosvitin^23,24^ and transported together with calcium ions in the form of calcium phosphate mineral vesicles. These mineral vesicles circulate in the bloodstream^22^ and play a key role in bone matrix mineralization^25^. Indeed, previous studies demonstrated that collagen sponges become mineralized when implanted on the CAM^26^. Yet, these prior experiments were performed with supraphysiological concentrations of bone morphogenetic protein-2 (BMP-2), which drives mineralization by stimulating MSC osteogenic differentiation regardless of matrix conditions. Moreover, fully understanding how the interplay between bone matrix, MSCs, and vascularization affects the type and location of mineral formed requires spatially resolved and correlative materials characterization that has not been achieved previously.

In this study, we take advantage of the CAM as a physiological bioreactor to reveal new insights into bone matrix mineralization. We have previously established protocols to decellularize bone matrices and selectively remove mineral^27^. This process preserves the micro- and nanostructure of bone’s organic matrix and allows for MSC seeding prior to CAM implantation^27,28^. Here, we leverage these protocols and implant decellularized bone matrices with and without mineral, and with and without MSCs, onto the CAM to study their independent and combined effects on matrix mineralization. Following explantation, we subjected the different tissues to advanced materials characterization, including correlative micro-computed tomography (µCT) and complementary analysis by Raman microscopy, scanning electron microscopy (SEM), and histochemical staining, enabling spatially controlled examination of matrix- and MSC-driven mechanisms involved in mineral deposition. Collectively, our results highlight that bone matrix mineral content regulates secondary, MSC-dependent mineralization in the CAM assay in an unexpected manner, and that the unique combination of tissue engineering strategies and high-resolution materials characterization enables insights not possible with traditional methods.

## Results

### Establishment of CAM bone matrix mineralization assay

To generate bone matrices with and without mineral, decellularized scaffolds were prepared following our previously established protocol^27^. Briefly, trabecular bone plugs were extracted from the subchondral bone of neonatal bovine femurs and subjected to post processing including sectioning, decellularization, and optional demineralization (Figure 1a-c). Successful scaffold decellularization was confirmed by the absence of nuclei in hematoxylin and eosin (H&E) stained cross-sections (Figure 1b). Additionally, μCT verified that mineral-containing bone scaffolds (termed ‘M-Bone’) exhibited mineralized trabeculae as expected, whereas demineralized bone scaffolds (termed ‘DM-Bone’) lacked contrast, indicating successful demineralization (Figure 1c). Importantly, the collagen structure of the trabeculae in DM-Bone remained intact after the demineralization process.

**Figure 1.**
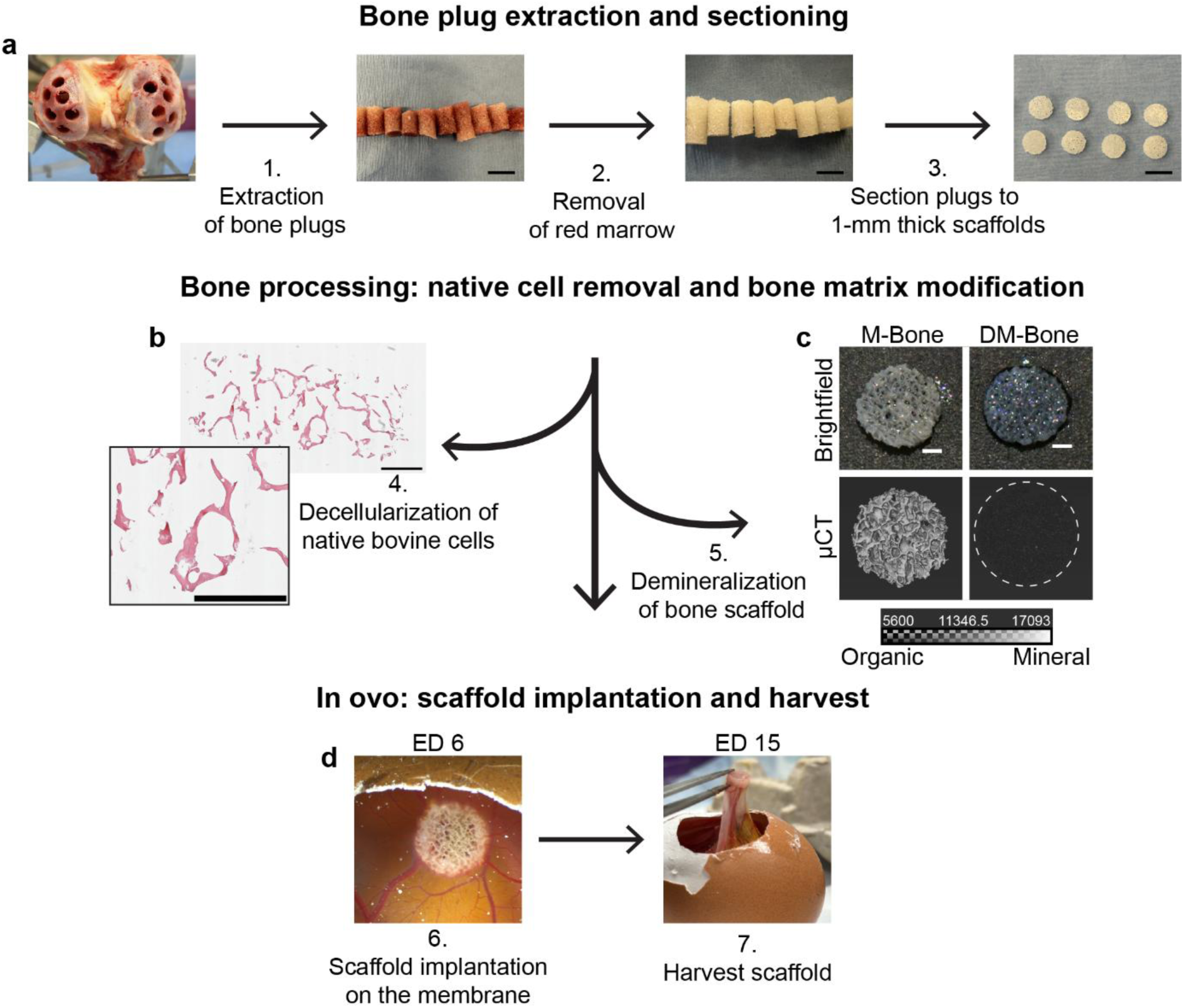
Overview of Experimental Setup. (a) Bone plugs were extracted from the subchondral bone of neonatal bovine femurs using a coring bit. Bone marrow was flushed out; bone plugs were sectioned into 6-mm x 1 mm cylindrical scaffolds. (b) Scaffolds were decellularized and (c) scanned with µCT to detect differences in mineral before and after demineralization. Mineral-containing bone scaffolds (M-Bone) showed high contrast indicating the presence of mineral whereas demineralized bone scaffolds (DM-Bone) showed low contrast, confirming that mineral was removed. (d) For *in ovo* experiments, scaffolds were placed on the CAM of a developing chick for 9 days (from embryonic day, ED, 6-15) and then harvested.

To analyze bone matrix mineralization using the CAM assay, we prepared eggs for scaffold implantation (Figure 1d). Embryogenesis was initiated by incubating eggs in a rocking-shelf incubator for 3 days simulating the natural movement during brooding. On embryonic day (ED) 3, a small window was created in the eggshell and sealed with semi-permeable tape and eggs were transferred to a stationary incubator. This approach permits the formation and expansion of the highly vascularized CAM while providing access for later scaffold implantation^29^. On ED 6, M-Bone and DM-Bone scaffolds were implanted onto the CAM of separate eggs, which were then re-sealed with tape for the remainder of the incubation period. Scaffolds were harvested after 9 days on ED 15; *i.e.*, prior to the development of immunocompetency^30,31^ and consistent with prior work^26,32,33^. Correlative imaging and analytical approaches were then combined to characterize mineralization and tissue formation in the M-Bone and DM-Bone scaffolds.

### CAM implantation leads to location-dependent matrix mineralization

We first used µCT to investigate how implantation onto the CAM affected mineralization of DM-Bone and M-Bone scaffolds (Figure 2a). Regardless of mineral content, implanted scaffolds became fully integrated in the CAM. Interestingly, mineralization of DM-Bone scaffolds only occurred on the side that faced the eggshell during incubation, while no mineral deposits were detected on the opposite side facing the embryo (Figure 2b). Although M-Bone scaffolds contained mineral both pre- and post-implantation, it was evident that scaffolds inoculated on the CAM had developed large mineral deposits on their periphery (Figure 2c). We quantified these observations by measuring the change in voxel intensity between spatially correlated μCT scans pre- and post-CAM implantation, which revealed that implantation onto the CAM increased mineralization for both scaffold conditions. Nevertheless, this increase was only statistically significant for the DM-Bone condition (Figure 2d,e) yielding similar results as for native bone.

**Figure 2.**
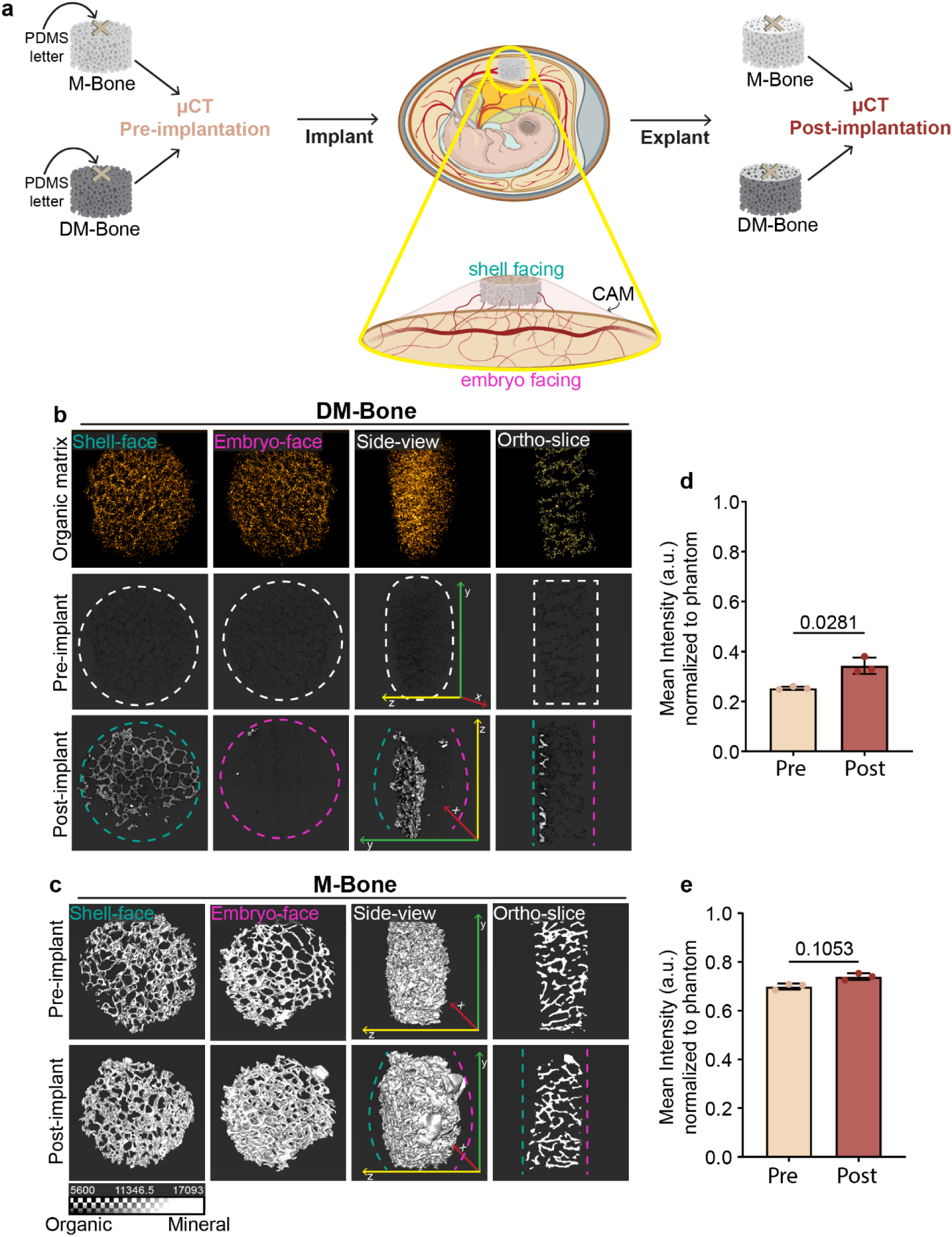
Implantation onto CAM promotes directional bone matrix mineralization. **a)** M-Bone and DM-Bone scaffolds were subjected to µCT before and after implantation onto the CAM to quantify the amount of newly mineralized matrix per scaffold. Each scaffold was labeled with a PDMS letter to maintain orientation during µCT scans and to help identify which side of the scaffold was facing the eggshell versus the embryo. **b, c)** Representative µCT images visualizing DM-Bone (b) and M-Bone (c) before (Pre-implant) and after (Post-implant) implantation onto the CAM. For visualization purposes only, all µCT images were adjusted to the same brightness and contrast level using ImageJ (FIJI). Axes shown in side view panels indicate scaffold orientation: red-x-axis, green-y-axis, and yellow-z-axis. Dashed lines indicate the location of the shell (teal) and embryo (magenta). **d, e)** Mineral content following CAM implantation was quantified from µCT scans of DM-Bone (d) and M-Bone (e) before and after CAM implantation. Statistical differences were determined using paired *t-*test with Wisconsin Sign Well test.

To resolve the spatial distribution of newly formed mineral deposits, voxel intensities from µCT datasets acquired prior to CAM implantation were subtracted from the corresponding post-implantation scans. The image output for DM-Bone scaffolds with (Figure S1b) and without (Figure 2b) prior subtraction showed that mineral only formed in shell-facing scaffold regions. Applying the same correlative approach to M-Bone scaffolds showed that although the net change in voxel intensity averaged over the whole bone plug was not significant (Figure 2e), there was nevertheless evidence of considerable flux with some regions experiencing new mineral formation, while in others mineral was removed (Figure S1d). Altogether, both DM-Bone and M-Bone showed most newly deposited mineral formed at the side that faced the shell, whereas the side closer to the embryo was more heterogeneous with some large newly-formed mineral clusters, as well as areas of mineral loss (Figure S1e,f). These findings suggest that organic bone matrix effectively guides new mineralization in the CAM assay, but only in areas adjacent to the shell. Our results also indicate that the CAM assay may be a valuable model to determine mechanisms involved in both new deposition and dissolution of mineralized bone matrix.

### Newly formed mineral in CAM assay consists of hydroxyapatite

To confirm the formation of new mineral and analyze its spatial distribution in more detail, histological cross-sections were stained with von Kossa and Alizarin Red to detect calcium phosphate^34^ and matrix-bound calcium ions^35^, respectively. Additionally, Picrosirius Red staining allowed visualizing the collagen-rich organic matrix of bone and infiltrated new tissue. Prior to implantation, DM-Bone scaffolds only stained positive with Picrosirius Red. Following CAM-implantation, von Kossa and Alizarin Red staining indicated mineralization occurring in shell-facing regions of DM-Bone scaffolds (Figure 3a). As expected, the fully mineralized M-Bone scaffolds stained uniformly with von Kossa and Alizarin Red, both before and after CAM implantation (Figure 3b). New tissue infiltrating the empty space between trabeculae was detected in both DM-Bone and M-Bone scaffolds, indicating that the bone plugs were fully integrated with the CAM.

**Figure 3.**
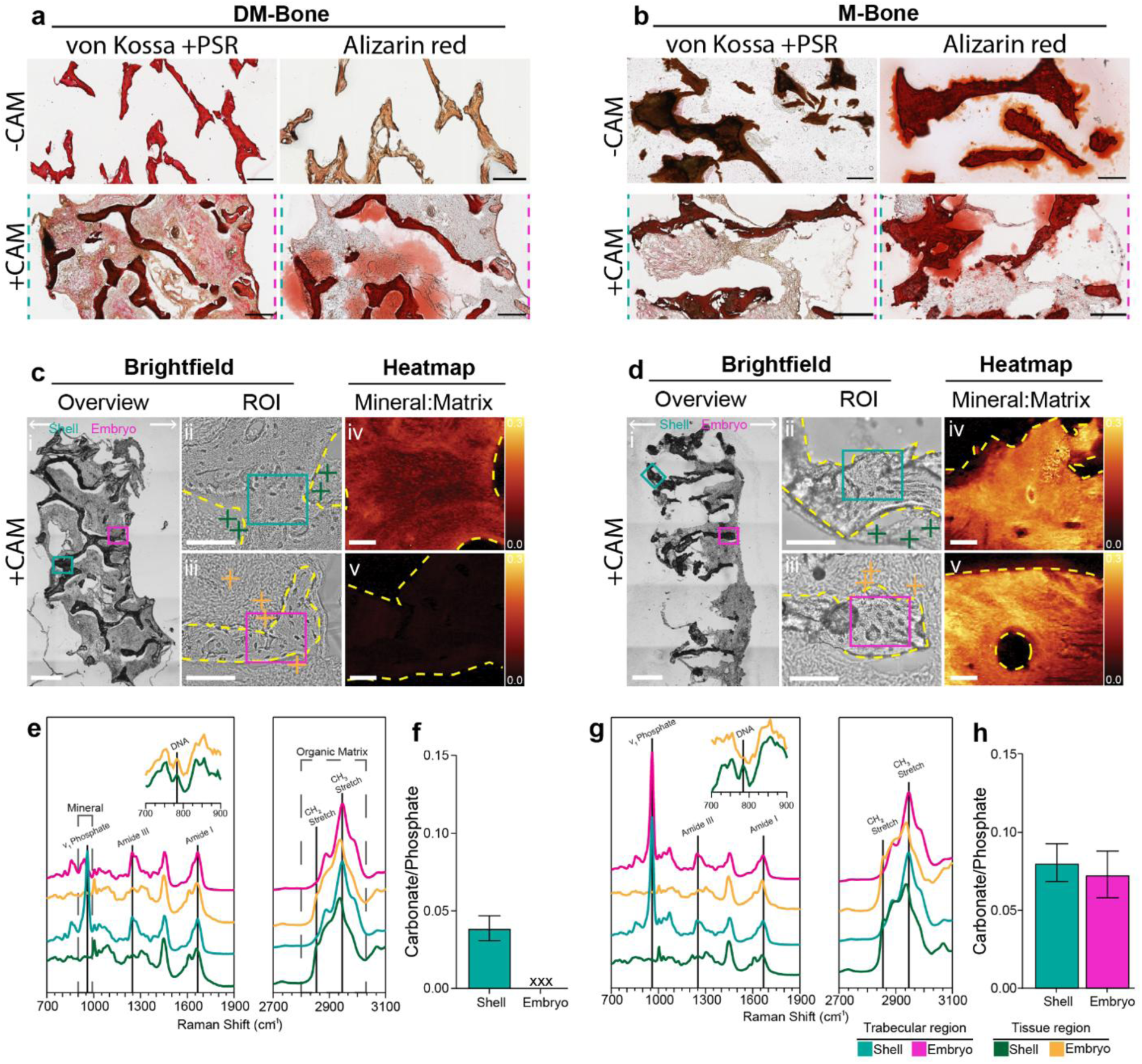
New mineral only formed in preexisting bone matrix and consists of hydroxyapatite. a, **b)** Von Kossa and Picrosirius Red (PSR) or Alizarin Red staining of DM-Bone (a) and M-Bone (b) cross-sections (scale bar: 200 µm). **c, d)** Raman microscopy: Unstained cross-sections (*i*) were used to identify ROIs for analysis by brightfield microscopy. Insets indicate ROIs of analyzed shell- (*ii*, teal) and embryo- (*iii*, magenta) facing regions; yellow dotted lines outline trabeculae; green and yellow crosses indicate newly formed tissue from which Raman spectra in e) and g) were collected (scale bar: 100 µm). Corresponding heatmaps of mineral to matrix ratio (*iv,v*) (scale bar: 20 µm). **e, g)** Raman spectra identified the chemical groups of the trabecular region (teal and magenta) and newly formed tissue (green and yellow) in the shell (teal and dark green) or embryo regions (magenta and yellow). **f, h)** Quantification of carbonate to phosphate ratio in trabeculae of shell and embryo regions.

To identify the composition of the mineral that formed on trabeculae during CAM implantation, confocal Raman maps were taken on unstained scaffold sections. Using brightfield microscopy, we first selected representative trabecular regions from shell- and embryo-facing interfaces of the scaffold explants for analysis (Figure 3c-h). Consistent with the mineralization pattern seen in µCT and histological staining, Raman spectra of DM-Bone scaffolds showed the presence of crystalline calcium phosphate peaks (ν_1_(PO_4_), 961 cm^-1^) characteristic of HA in shell-, but not embryo-facing, regions of the scaffold (Figure 3e). The typical amide III peak complex (1250 cm^-1^) and strong proline (860, 923 cm^-1^) and hydroxyproline (876, 943 cm^-1^) peaks indicative of collagen I were detected in the trabecular matrix in both regions. A comparable mineral and organic matrix composition was likewise found in M-bone scaffolds in both shell- and embryo-facing regions (Figure 3g).

The degree of mineralization was spatially quantified with Raman maps by comparing the mineral:matrix peak intensity ratio in both scaffold types, revealing high contrast indicative of mineral in the shell-, but not the embryo-facing region of DM scaffolds (Figure 3c (*iv-v*)) whereas M-Bone displayed high mineral contrast in both regions (Figure 3d (*iv-v*)). Consistent with the formation of newer, less mature mineral, the carbonate content of HA (quantified through the carbonate:phosphate peak intensity ratio) of shell-facing regions in DM-Bone scaffolds (Figure 3f) was lower than in both the embryo- and shell-facing regions of M-Bone scaffolds (Figure 3h). Since the carbonate:phosphate parameter is specific to HA, it could not be calculated for the embryo-facing region of DM-Bone where no mineral was found. Raman spectra of the new tissue that had infiltrated the space between trabeculae suggested that it consisted of non-collagenous proteins, lipid, and nucleic acids consistent with cells, but did not contain any collagen or mineral regardless of scaffold type (Figure 3e,g). Collectively, these results suggest that bone matrix mineralization in the CAM assay yields a physiologically relevant mineral phase consistent with newly-formed bone-like HA that is less mature than preexisting native mineral in M-Bone scaffolds, and that the new mineral formed only in bone matrix (*i.e.*, trabeculae) but not in newly infiltrated tissue.

### Formation of new mineral negatively correlates with density of infiltrated host cells

As mineralization is often thought to be a cell-driven process, we next evaluated how host cells invaded DM-Bone and M-Bone scaffolds implanted onto the CAM. Using image analysis of H&E-stained histological cross-sections, we identified that while native cells invaded equally well into both scaffold systems, the density of infiltrated stromal cells decreased in the direction of the eggshell in both cases (Figure 4a-d). Quantification of cell density confirmed that the infiltrated new tissue was more abundant in embryo-facing scaffold regions relative to shell-facing regions, and therefore, opposite to our matrix mineralization findings. Indeed, a correlation analysis likewise showed that mineralization of DM-Bone negatively correlates with cell density (Figure 4e).

**Figure 4.**
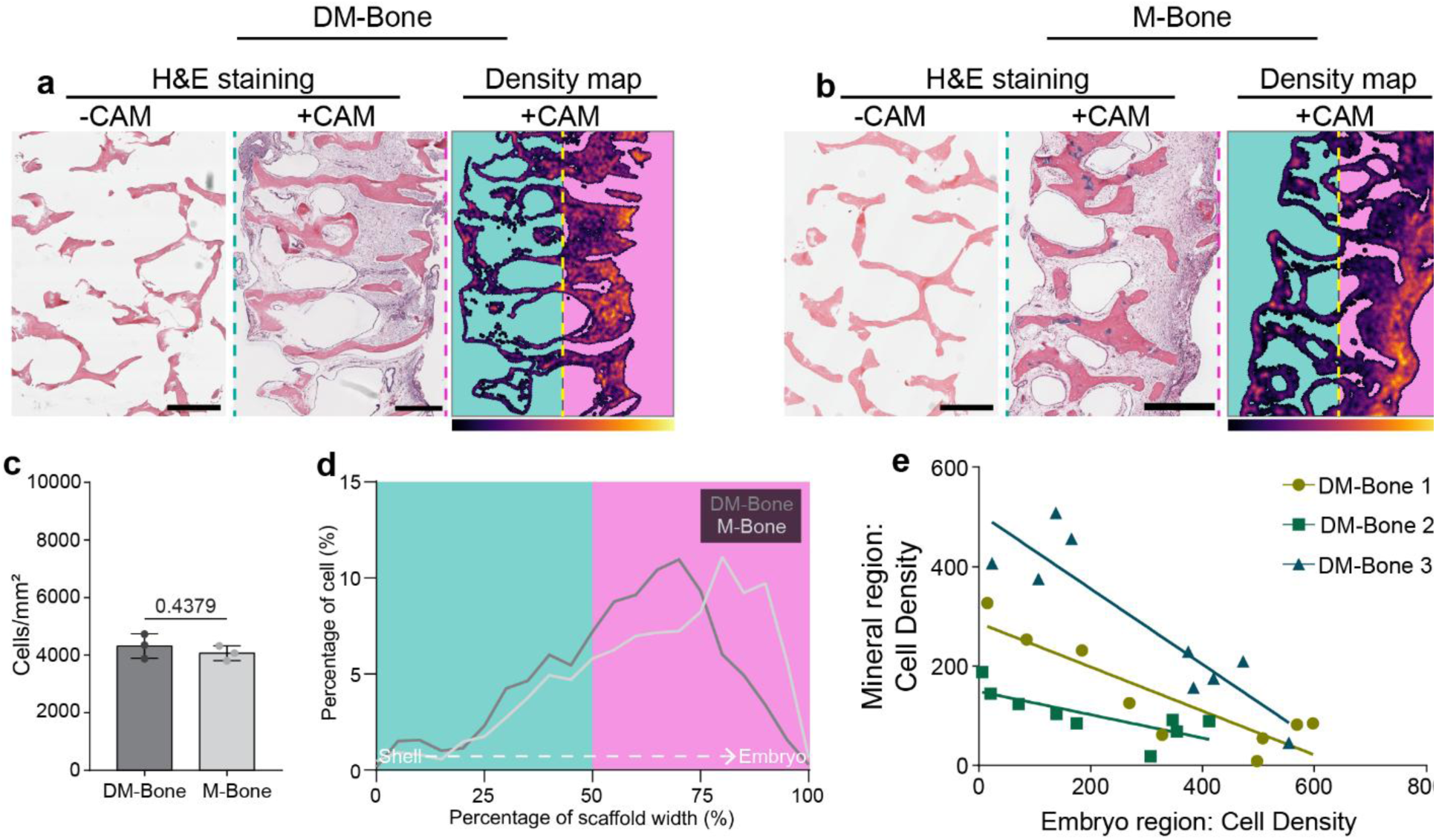
Density of infiltrated cells decreases with increasing distance from CAM. a, **b)** Representative H&E images of scaffolds before and after CAM implantation and corresponding cell density maps of DM-Bone (a) and M-Bone (b) as quantified by image analysis of cell nuclei (scale bar: 500 µm). **c)** Quantification of cell density per scaffold area (DM-Bone: n=3, M-Bone: n=3). **d)** Histogram of cell distribution (%) across the scaffold thickness from the shell side to the embryo side. **e)** Correlation analysis (DM-Bone 1: *r*(7) = - 0.72, *p =* 0.037; DM-Bone 2: *r*(7) = -0.78, *p* = 0.017; DM-Bone 3: *r*(7) = -0.80, *p* = 0.014) of native chick cell density found in mineral and embryo region.

### Establishing the presence of MSCs within scaffold system

To more directly evaluate how the presence of stromal cells affects mineralization in the CAM model, we seeded human bone marrow-derived mesenchymal stem cells (MSCs) into the scaffolds prior to implantation (Figure S2a). Using a DNA assay we ensured equal seeding density of MSCs (Figure S2b) in both scaffold types prior to implantation. Fluorescent imaging of transverse sections additionally confirmed that MSCs adhered to trabecular surfaces and were homogeneously distributed throughout both scaffold types (Figure 5a, S2c). Following implantation, human MSCs were also located in the newly infiltrated tissue as revealed by immunofluorescent staining of human vimentin. Interestingly, post-implant DM-Bone contained human MSCs only on the shell-facing interface (Figure 5a), whereas in M-Bone scaffolds human MSCs were present throughout (Figure S2c). We validated these qualitative observations by quantifying the distribution of fluorescence intensity of human MSCs before and after implantation (Figure 5b, S2d), which confirmed their retention in both systems but different spatial distribution in DM-Bone vs. M-Bone scaffolds.

**Figure 5.**
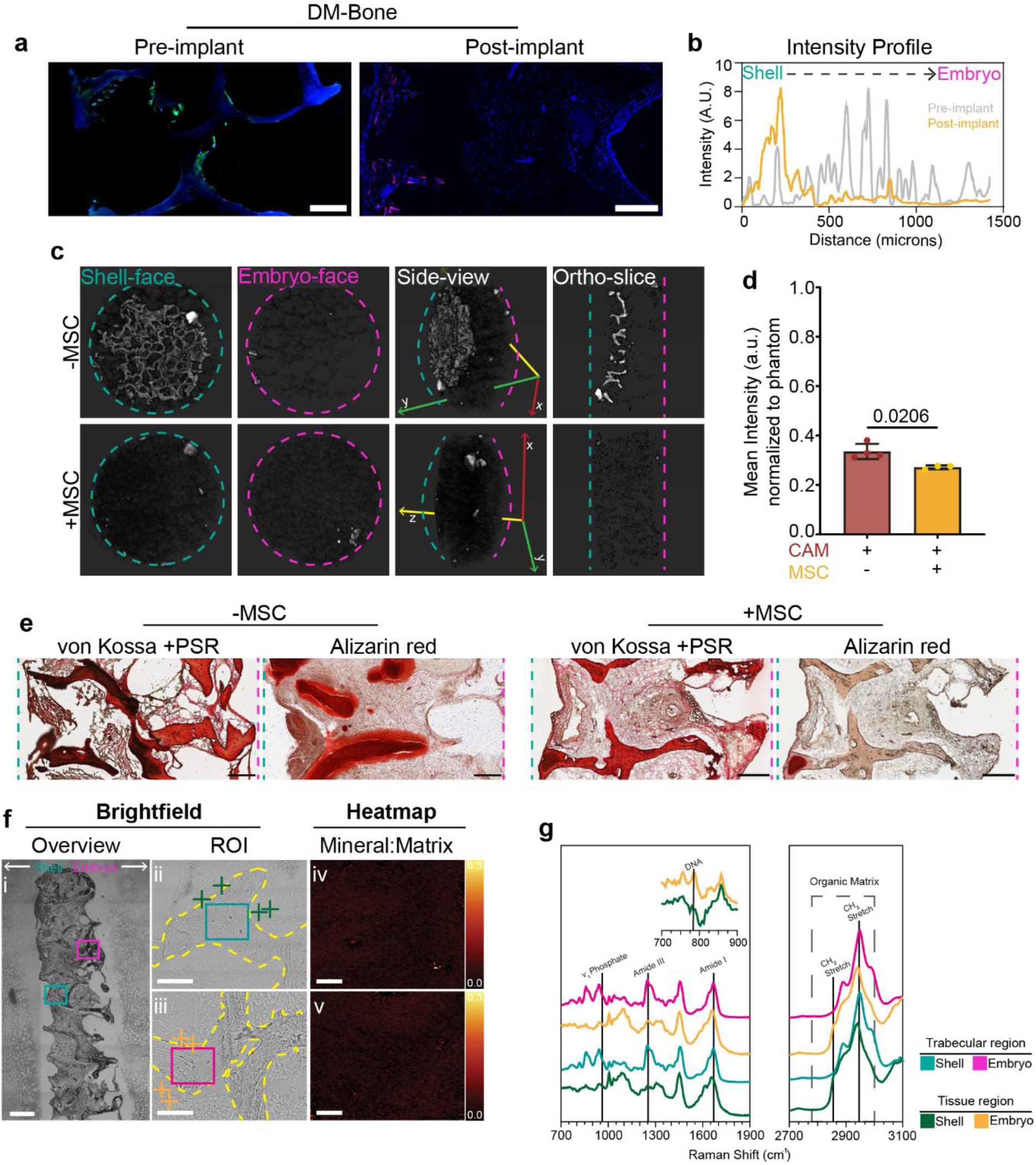
Pre-seeding scaffolds with MSCs prior to CAM inhibits mineralization. **a)** Fluorescence images of MSCs seeded into DM-Bone scaffolds pre- and post-implant (Pre-implant: Phalloidin (green) and DAPI (blue), Post-implant: Human vimentin (magenta) and DAPI (blue)) (scale bar: 200 µm). **b)** MSC presence pre- and post-implant was quantified by measuring the intensity profile of the phalloidin or vimentin fluorescence intensities across sample sections. **c)** Mineral assessment of DM-Bone +/- MSCs via µCT. **d)** Mineral density of DM-Bone +/- MSCs measured from µCT images. **e)** DM-Bone +/- MCSs stained with von Kossa and PSR or Alizarin Red (scale bar: 200 µm). **f)** Raman microscopy: overview brightfield image (*i*) (scale bar: 500 µm), region of interest (*ii*) (scale bar: 100 µm), and mineral to matrix ratio heatmap (*iii*) (scale bar: 20 µm). **g)** Raman spectra identified only organic matrix.

### MSCs interfere with bone matrix mineralization

Since MSCs can facilitate bone formation^36^, we returned to µCT to examine how pre-seeding with MSCs affects mineral deposition of scaffolds implanted on the CAM. As expected, DM-Bone and M-Bone without MSCs showed similar mineralization patterns as described above (Figure 5c, S3a). However, in the presence of MSCs, mineral deposition on DM-Bone was reduced, and M-Bone did not have the large mineral clusters seen in the MSC-free CAM assay. Further quantification of µCT voxel intensity revealed a decrease in mean intensity for both scaffold systems when they were seeded with MSCs prior to implantation (Figure 5d, S3b).

Consistent with the µCT analysis, MSC-seeded DM-Bone scaffolds exhibited dramatically reduced von Kossa and Alizarin Red staining after CAM implantation as compared to DM-Bone scaffolds that were not seeded with MSCs (Figure 5e). Accordingly, Raman microscopy of MSC-seeded DM-Bone sections (Figure 5f-g) showed no contrast in the heatmaps of either the shell or embryo regions (Figure 5f (*iv-v*)), and corresponding spectra (Figure 5g) showed the collagen typical of DM-Bone but no sign of mineral. No changes in histochemical staining or Raman signature were observed in MSC-seeded M-Bone scaffolds compared to their controls (M-Bone without MSC) (Figure S3c-f). Altogether, these findings suggest that MSCs inhibited the deposition of new mineral but did not affect the chemical composition of any pre-existing mineral.

### Osteogenic induction of MSCs does not restore bone matrix mineralization on DM-Bone

To further investigate MSC’s role during matrix mineralization in the CAM, we hypothesized that promoting MSC differentiation toward osteoblast-like cells would restore mineral deposition. Thus, MSCs were osteogenically differentiated for one week prior to seeding on DM-Bone scaffolds. To validate the differentiation process, we performed an assay for alkaline phosphatase (ALP), an early marker of osteogenic differentiation. Despite increased ALP activity following one week differentiation (Figure S4a) indicating that MSCs had been pushed toward an osteoblast-like phenotype before seeding them on the scaffolds and implanting them onto the CAM, no mineralization of DM-Bone was detected by von Kossa and Alizarin Red staining (Figure S4b). These results suggest that the presence of MSCs, regardless of osteogenic priming, negatively interferes with bone matrix mineralization in the CAM assay.

### Microvessel density is increased at the shell-facing scaffold surface

Because we observed spatial differences in DM-Bone scaffold mineralization and because others have reported that the CAM vasculature mediates mineralization by transporting calcium and phosphate ions in the form of calcium phosphate mineral vesicles ^23,24^, we next speculated that spatial differences in bone matrix mineralization of DM-Bone scaffolds are due to differences in microvessel integration. Interestingly, brightfield imaging suggested that many small capillaries infiltrated the shell-facing interface of scaffolds (Figure 6a(*i*)) whereas larger vessels were visible on the embryo-facing interface (Figure 6a(*ii*)), which we further validated using second harmonic generation (SHG) and fluorescence imaging following perfusion with Texas Red-dextran (Figure 6a(*iii*)). These macroscopic differences prompted us to quantify microvessel density of shell- and embryo-facing scaffold regions in more detail (Figure 6b). Indeed, microvessel density on the shell- vs. embryo-facing scaffold interface was increased (Figure 6c), a trend that we additionally validated by quantification of lectin-stained blood vessels (Figure 6d-e). These results indicate that increased bone matrix mineralization of DM-Bone scaffolds in the CAM assay correlates with elevated microvessel density, but reduced cell infiltration.

**Figure 6.**
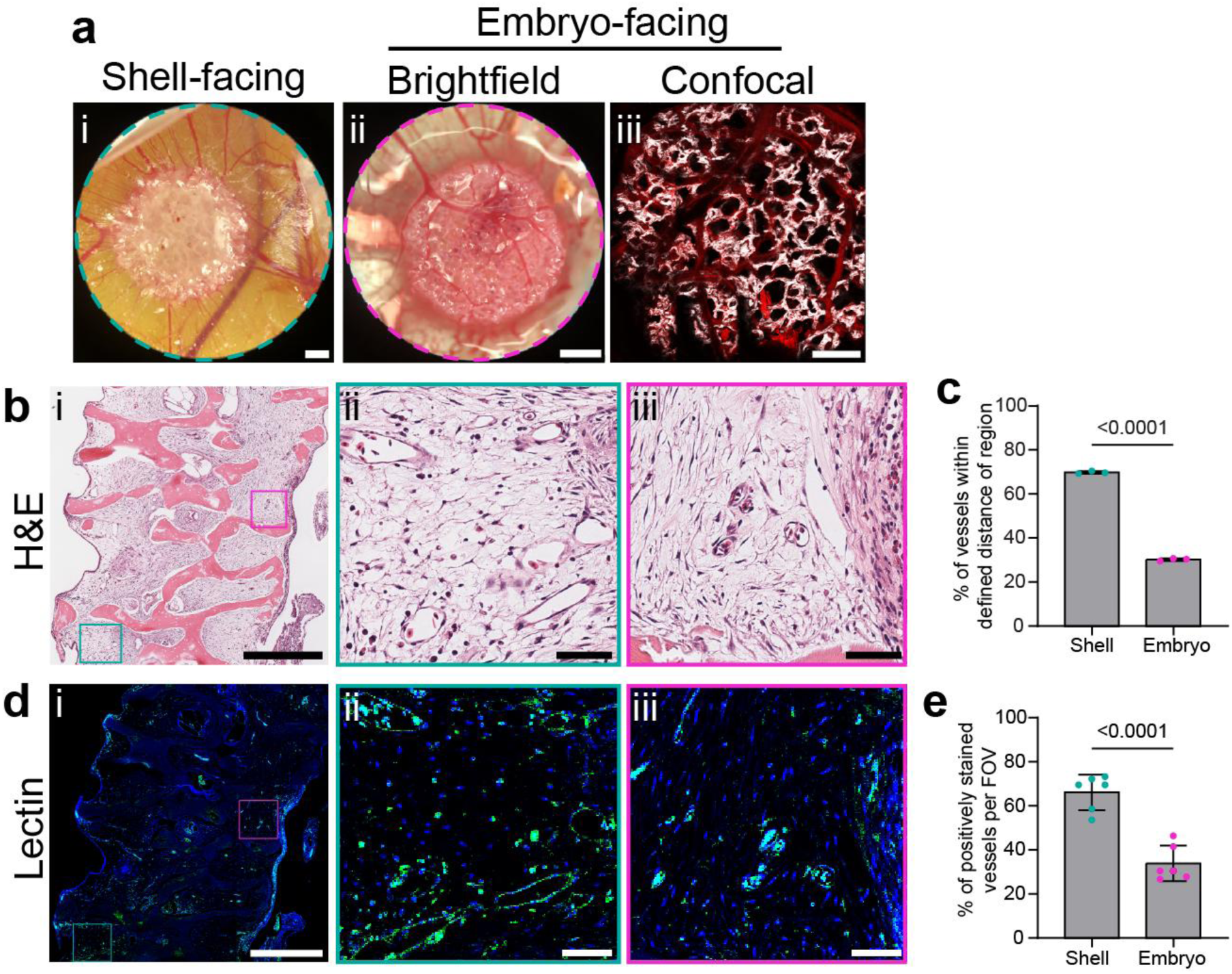
Microvessel density is greater on shell-facing interface of DM-Bone. **a)** Stereoscopic images of shell-facing (*i*) (scale bar: 1 mm) and embryo-facing (*ii*) side of DM-Bone scaffolds implanted on CAM (scale bar: 5 mm). (*iii*) Representative SHG and Texas Red fluorescence image of the embryonic side of DM-Bone scaffold showing matrix architecture and perfused blood vessels (scale bar: 1 mm). **b,d)** H&E staining and lectin staining of histological cross-sections visualizing microvessels within DM-Bone (scale bar: 500 µm) (inset scale bar: 50 µm). **c)** Quantification of microvessels in shell or embryo-facing region of H&E staining. **e)** Quantification of lectin-stained microvessels per FOV.

### Bone matrix mineralization in the CAM assay depends on microvessel integration

To test the role of microvessel-mediated rather than diffusion-dependent transport of mineral precursors in the mineralization of DM-Bone scaffolds more directly, we embedded DM-Bone scaffolds into 7% agarose gels. Previous studies have shown that adjusting agarose concentration can be used to control hydrogel porosity to allow for diffusion of small and large molecules, while preventing cellular infiltration (Figure 7)^37–39^. Indeed, incubation of agarose-embedded DM-Bone scaffolds in a polymer-induced liquid precursor (PILP) mineralization solution^40^ containing calcium, phosphate, and polyaspartic acid *in vitro* confirmed that this approach allowed for diffusion of mineral-specific biopolymers and soluble ions into the agarose gel and consequently resulted in scaffold mineralization - although at slightly reduced levels as compared to non-encapsulated controls (Figure 7a). Next, we implanted agarose-embedded scaffolds onto the CAM which led to integration of the agarose-embedded DM-Bone scaffold within the CAM and submersion in allantoic fluid. However, this protocol also blocked blood vessel and stromal cell infiltration and, simultaneously, prevented mineralization (Figure 7b). Consequently, these results suggest that blood vessel infiltration and transport of circulating factors is essential in mediating mineralization of DM-Bone scaffolds in the CAM assay.

**Figure 7.**
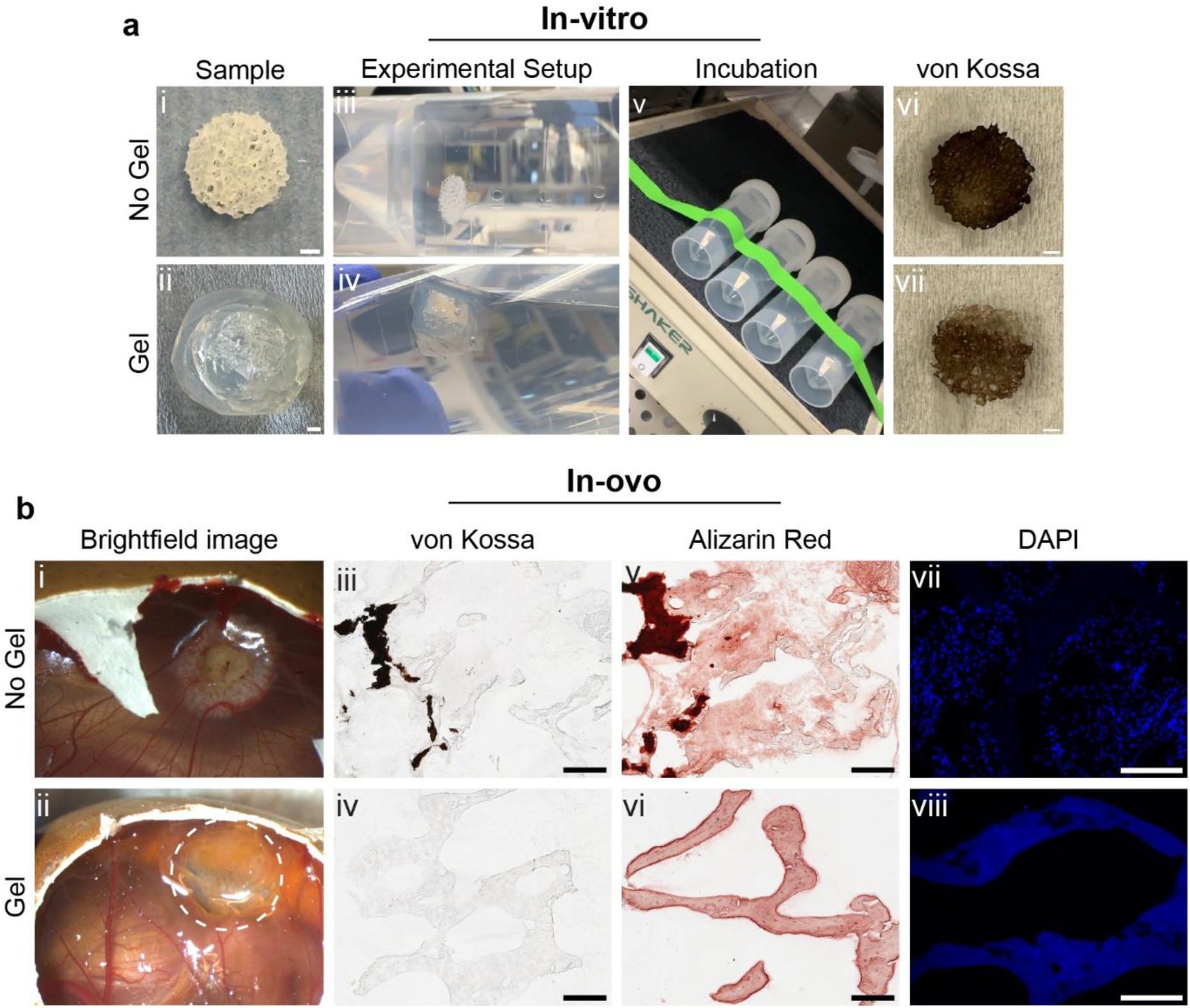
Agarose encapsulation allows diffusion of soluble factors while blocking cell invasion. **a)** Representative image of control (*i*) and agarose-encapsulated DM-Bone scaffold (*ii*) before and after in vitro mineralization using the PILP process (scale bar: 1mm). Mineralization of scaffolds incubated (*iii,iv*) in mineralization solution (*v*) was confirmed by von Kossa staining (*vi,vii*). **b)** *In ovo* mineralization of DM-Bone scaffold (*i*) and agarose-encapsulated DM-Bone scaffold (*ii*). Cryosections of scaffolds following CAM implantation stained with von Kossa (*iii,iv*), Alizarin red (*v,vi*), or DAPI (*vii,viii*) (scale bar: 200 µm).

### Matrix remodeling induced by MSC interferes with vascular integration

As our results suggest a positive correlation between microvessel infiltration and DM-Bone scaffolds mineralization, while the presence of MSCs was negatively correlated with mineralization, we hypothesized that MSCs disrupt bone matrix mineralization by altering microvessel formation. Quantification of vessel density in histological cross-sections (Figure 8a-c) revealed that the presence of MSCs significantly decreased total vessel density (Figure 8b) and vessel density on the shell interface, while increasing vessel density on the embryo interface (Figure 8c). We have previously demonstrated that MSCs seeded on collagen type I, the main organic bone matrix protein, induce a pro-inflammatory environment characterized by profibrotic matrix accumulation and contractility^14^, the latter of which has been shown to exert anti-angiogenic effects^41,42^.

**Figure 8.**
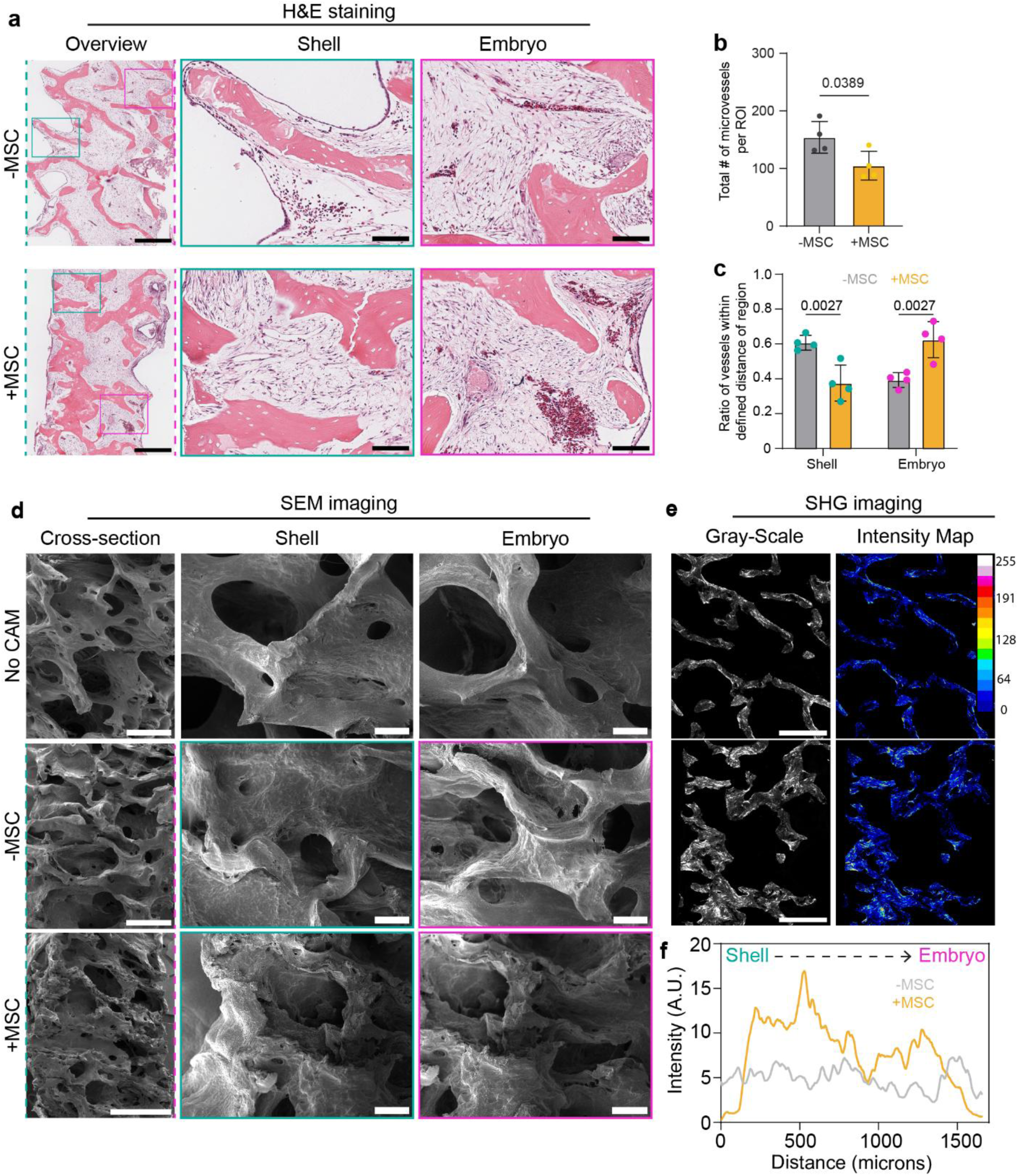
MSCs alter vessel density and trabecular architecture of scaffolds. **a)** Representative H&E staining of DM-Bone +/- MSC (scale bar: 100 µm, 500 µm). **b)** Total microvessel density within DM-Bone as quantified by image analysis of H&E-stained cross-sections. **c)** Quantification of microvessel density in shell or embryo-facing region of DM-Bone +/- MSC. **d)** Scanning electron microscopy of DM-Bone cross-section; no CAM implantation control and following CAM implantation +/- MSCs (scale bar: 100 µm, 1 mm). Teal dash lines indicate shell-facing and magenta dash lines indicate embryo. **e)** Second harmonic generation imaging of histological samples (scale bar: 500 µm). **f)** Quantification of collagen intensity profile from shell to embryo of DM-Bone +/- MSC following CAM implantation.

Given the pro-inflammatory and matrix remodeling responses of MSCs and that collagen nanostructure and gap zones are critical regulators of bone-like intrafibrillar mineralization^43–45^, we next examined DM-Bone scaffolds seeded with MSCs using SEM and SHG (Figure 8d,e). Interestingly, SEM images revealed that trabeculae appeared more porous, partially degraded and more compact in the presence vs. the absence of MSCs following CAM implantation (Figure 8d). SHG analysis further showed increased collagen density at the shell interface (Figure 8f); i.e., the location where MSCs were identified after DM-Bone scaffolds were retrieved from the CAM (Figure 5a). Altogether, these findings suggest that mineralization of DM-Bone scaffolds in the CAM depends on blood vessel infiltration and maintenance of bone matrix micro- and nanostructure, but that MSCs may interfere with this process by proteolytic bone matrix remodeling.

## Discussion

The CAM assay is increasingly explored as a tool to study bone formation and regeneration^26,32,33,46^, but the mechanisms by which mineral deposition is regulated within the CAM remain unclear. Improving our understanding of these mechanisms is critical because matrix mineralization governs bone integrity and may inform future studies exploring bone matrix remodeling in health and disease. Using a combination of bone-derived extracellular matrices and high-resolution analytical techniques to characterize matrix mineralization, we show that the microstructural properties of bone trabeculae regulate new mineral deposition and that direct access to the vasculature and thus, circulating factors such as mineral vesicles is essential for mineralization to occur in the CAM assay.

Previous studies investigating matrix mineralization using the CAM assay have used polymer-based scaffolds^26,46^ and did not report a similar degree of mineralization as we have found with our bone matrices, underscoring the importance of matrix composition and organization for bone-like matrix mineralization. Interestingly, the CAM itself is composed of similar extracellular matrix proteins as our scaffolds (e.g., collagen, fibronectin, laminin, and other proteins^47^), although, while we expected to observe mineralization on the membrane itself, we only identified HA deposition on the scaffolds’ trabeculae. Thus, DM-Bone and M-Bone served as an effective template, structured to support subsequent mineralization.

A previous study inducing defects within human bone cylinders to study bone regeneration in the CAM demonstrated ectopic mineral deposition and bone resorption^26^. Consistent with these results, M-Bone showed new mineral clusters on the periphery of the scaffold whereas mineral degradation occurred on the embryo-facing surface. Because our results showed void regions and tissue formation due to host cell infiltration, we attribute mineral loss to cell-mediated resorption. Indeed, previous research has suggested that chorion cells contained in the CAM secrete acid that dissolves calcium carbonate from the eggshell, endocytose calcium ions, and transport the calcium ions to the endoderm of the membrane in a vesicle-dependent manner^48^. Therefore, the changes in M-Bone new mineral deposition and removal could be due to chorion cells, a hypothesis that will need to be verified in future studies.

We detected a surprising gradient of bone matrix mineralization in DM-Bone scaffolds where shell-facing regions of the scaffolds were more mineralized than their embryo-facing counterparts, and this increase correlated with decreased cell density and increased microvessel integration. The elevated cell density at the embryo-facing interface is likely because the distance for host cells to infiltrate embryo-facing scaffold regions is much shorter than shell-facing scaffold regions. Hence, host cells have more time to proteolytically remodel bone matrix before effective vascular ingrowth and provision of mineral precursors. Collectively, these results suggest that vascular-mediated delivery of circulating mineralization factors constitutes the rate-limiting step during organic bone matrix mineralization and must be tightly coordinated with cell-seeding and/or host cell invasion to prevent excessive remodeling or degradation of the matrix prior to effective mineral deposition and stabilization.

Our results show selectively blocking cell infiltration through agarose encapsulation or pre-seeding DM-Bone scaffolds with MSCs inhibited *in ovo* mineralization, and vascular integration was greatly limited in both cases^49–51^. This is notable, given that the vasculature serves as a primary transport system for essential molecules^49^. Prior studies have demonstrated that, during chick embryonic bone development, soluble ion concentrations greatly exceed the amounts required for bone formation, indicating an abundant and readily available calcium ion supply^48^. Thus, high calcium influx could potentially be a critical factor in driving new mineral formation in our experiments. Indeed, prior *ex ovo* experiments have shown improved bone development when removing chick embryos from the shell^50^ by adding crushed shell (i.e. shell dust) to the vascularized membrane to supplement calcium^51,52^. Nonetheless, these observations indicate that sustained calcium exposure, coupled with vascular-mediated transport^49^, is critical for bone formation and likely underlies mineral deposition in our experiments. While further studies are needed to determine the exact mechanisms, these studies suggest that matrix mineralization within the CAM relies primarily on the circulation of calcium and phosphate ions through the vascular network.

Altogether, our results emphasize the diverse use of the vascular bed within the CAM. For example, the CAM assay has been a long-favored system for studying angiogenesis or drug screening during tumor growth^53–55^, and has been largely overlooked as a model for other biological processes (i.e. matrix mineralization). Additionally, our results highlight the importance of bone matrix organization in directing and supporting vasculature integration and cellular infiltration during the mineralization process. These findings transform the CAM assay into a versatile and biologically rich model for bone matrix mineralization, enabling a cell-free, rapid biomineralization without the need of additional growth factors. Lastly, unlike prior studies that have focused primarily on disease-specific bone regeneration within the CAM, our work investigated spatiotemporal dynamics induced by the CAM and its implications in cell-mediated bone matrix remodeling. While the CAM assay is not a definitive replacement for mouse models, it offers an inexpensive, high-throughput, and physiologically relevant system to help study key parameters of mineralization processes, including mass transport, vascularization, matrix organization, and cellular remodeling.

## Materials and Methods

### Scaffold fabrication

6-mm diameter bone plugs were extracted from neonatal bovine femurs (Copper City Meats: Rome, NY). The red marrow was flushed out under a high velocity stream of deionized water and bone plugs were subsequently sectioned into ∼1mm thick scaffolds. For decellularization, these scaffolds were incubated in extraction buffer containing 20 mM NaOH and 0.5% Triton X-100 in phosphate buffered saline (PBS) and then in 20 U/ml DNAse I to remove any lingering DNA fragments. For demineralization, bone scaffolds were placed in a 9.5% ethylenediaminetetraacetic acid (EDTA) (G Biosciences) solution overnight at 4 °C with low stirring, followed by multiple wash steps of PBS and deionized water. Prior to CAM implantation, scaffolds were sterilized with 70% ethanol (EtOH) for 10 minutes, followed by multiple wash steps with PBS.

### Chick embryo preparation

Fertilized chick eggs (Westwind Farms; Ovid, NY) were stored at 4 °C. At embryonic day (ED) 0, chick eggs were transferred to an incubator set between 35 to 40 °C with rotating shelves to prevent attachment of the yolk to the eggshell. At ED 3, a small window within the eggshell was created to enable future inoculation. This window was then sealed with semi-permeable tape (Saniderm) to control temperature and oxygen tension. The chick egg was then transferred to a stationary incubator (Genesis Hova-Bator). At ED 6, the semi-permeable tape was gently opened, scaffolds were inoculated, and a fresh semi-permeable tape was replaced to close the window. At ED 15, chick embryos were euthanized and scaffolds were harvested.

### Micro-computed tomography (µCT) of scaffolds

Individual scaffolds were placed in microcentrifuge tubes with 70% EtOH following fixation and were then aligned in 50 mL tubes for imaging. Image scans were collected using a Bruker SkyScan 1276 at 10 microns/pixel. A phantom cylinder image was collected simultaneously for image calibration and normalization. 3D virtual images were rendered using parameters such as volume and intensity measured by Avizo software (Thermo Fisher Scientific). Briefly, a segmentation of individual scaffolds was performed manually using the wand tool, creating a binary image label. The binary label was used as a mask on the original dataset to generate an image with identical voxel dimensions and spatial properties. Scaffold images and quantifications were created using tools like volume rendering, ortho slice, global analysis, or as indicated otherwise.

To maintain scaffold orientation for µCT, polydimethylsiloxane (PDMS) (Dow Corning) was prepared via a kit mixture at a ratio of 10:1 and cured overnight at 60 °C. Utilizing a sharp edge, letters were carved out of the PDMS mold. 6-mm biopsy punches (Integra Miltex) were used to cut out circular-shaped blue foam biopsy pads (VWR International) to be used as dividers. In 2 mL centrifuge tubes, scaffolds were submerged in 70% EtOH and arranged in the following order to keep track of their orientation: blue foam pads, scaffold, PDMS letter, and repeated. µCT captured scans were collected as previously described. Following imaging, scaffolds underwent PBS washes in preparation of CAM implantation. Post-harvest, scaffolds underwent fixation using 4% paraformaldehyde (PFA) and PBS washes. Utilizing the PDMS letters to distinguish eggshell and embryo interface, scaffolds were placed in 2 mL centrifuge tubes as described above for µCT scans.

### Avizo arithmetic computation

Segmentations of scaffolds pre- and post CAM implantation were aligned utilizing the transformer tool prior to registering the image. Applying a bounding box to both images, lattice axes were aligned to ensure both images have the same dimensions. Using the subtract image tool, the intensity of pre-CAM implantation images was subtracted from their corresponding intensity of post-CAM implantation images to generate the final image.

### Cryoembedding and sectioning for bone matrix characterization

Following fixation, scaffolds were dehydrated via a sucrose (Sigma-Aldrich) gradient for subsequent embedding in OCT. A cryotome (Thermo Scientific Cryostat Microm, HM 550) was then used to section 10 µm-thick slices. Samples were mounted onto glass slides (Globe Scientific).

Mineral deposition on cryosections was assessed using von Kossa and Alizarin Red S staining. Silver nitrate (Sigma Aldrich, S8157-10G) powder was dissolved in deionized (DI) water to prepare a 1% (w/v) solution. Sodium thiosulfate (Sigma Aldrich, S7026-250G) was similarly dissolved in DI water to obtain a 2.5% (w/v) solution. Samples were counterstained with Picrosirius Red (Electron Microscopy Sciences, 26357-02). For Alizarin Red staining, Alizarin Red (VWR, 97062-616) powder was dissolved in DI water and pH-adjusted to obtain a 40 mM solution at 4.1.

### Confocal Raman Microscopy and Raman Map Analysis

Using a WITec alpha300R confocal Raman microscope (WITec GmbH, Ulm, Germany), hyperspectral Raman maps were acquired from representative regions on the shell- and embryo-facing sides of 10 μm thick cross-sections immersed in PBS. All maps were acquired with a Zeiss W Plan-Apo 63×/1.0 water-immersion objective at 1 μm pixel size and 0.1 s integration time, with an excitation wavelength of 532 nm and a 300 lines/mm diffraction grating. Calibration spectra were recorded with a synthetic hydroxyapatite standard. WITec Project 5 software was used for all Raman map analysis. Spectra were baseline-corrected using a shape size of 500, and cosmic rays were removed manually where present. Each map was then masked to separate the pixels belonging to the trabecular and tissue regions, and noise reduction was performed by binning adjacent pixels. The identities of major mineral and organic components were confirmed by their characteristic Raman signatures (Table S1). Bone composition parameters were quantified from the spectral abundances of crystalline calcium phosphate mineral, carbonate ion lattice substitutions, and organic matrix by integrating the intensities of the ν_1_(PO_4_^3-^) and ν_1_(CO_3_^2-^) bands and the CH_x_ stretching peak complex, respectively, over the ranges listed in Table S2. The mineral:matrix and carbonate:phosphate ratios were then calculated by dividing the ν_1_(PO_4_^3-^) and CH_x_ stretch, and the ν_1_(CO_3_^2-^) and ν_1_(PO_4_^3-^) peak areas, respectively.

### Histological staining and imaging

Harvested scaffolds were submitted to Cornell’s Pathology and Laboratory Core for paraffin embedding, sectioning and H&E staining. Unstained paraffin sections were used for immunostaining with rabbit anti-human vimentin (Thermo Scientific, MA5-16409).

Brightfield images were captured with a ×40 air objective using an Aperio ScanScope CS2 (Leica Biosystems). To quantify cell recruitment, H&E images were uploaded to QuPath v0.5.0 and the cross-sectional scaffold area was manually annotated to include cells and exclude trabeculae. Each data point represents the average cell density (cells per mm^2^), calculated from sister sections of the same sample. Cell density maps were computed using Qupath’s train pixel classifier to distinguish two regions: tissue and trabeculae. Based on the classifications created, the density map feature was used to generate hotspots in the tissue region. Each pixel relates to the density of “objects” within a defined region.

### Vasculature characterization

Adequately sized blood vessels were perfused with lysine fixable Texas-Red Dextran (Thermo Scientific, D3328) dye through the vascular bed using a fine tip capillary needle. Following incubation, the membrane was then perforated using a cauterizing tool (Bovie Medical) to seal blood vessels and minimize blood leakage during explantation.

For fluorescence imaging, fluorescein-lectin (Thermo Scientific, L32479) was added to samples at a dilution of 1:100. DAPI (Thermo Scientific, D1306) was used as a counterstain to visualize nuclei. Following staining, Sudan Black (Sigma-Aldrich, 199664-25G) was added at a concentration of 10 mg/mL to minimize autofluorescence.

### Human MSC culture

MSCs (RoosterBio) were cultured in RoosterNourish-MSC media until 80% confluency. MSCs were seeded on scaffolds (500,000 cells/mL) in 50 µL of Dulbecco’s Modified Eagle Medium (DMEM), 10% FBS and 1% P/S.

### Correlative imaging approaches

To visualize trabecular structure, scaffolds were imaged using scanning electron microscopy (SEM) (Mira3 LM, Tescan). Briefly, scaffolds were fixed in 2% glutaraldehyde in sodium cacodylate buffer, followed by dehydration using a serial ethanol gradient, and imaged at 5 kV. Additional second harmonic generation (SHG) images of the collagen microstructure were captured with a ×32 water immersion objective using a LSM880 Inverted Confocal Multiphoton microscope (Zeiss) and analyzed with ImageJ.

### Statistical analysis

Unless otherwise indicated, results are presented as the mean and standard deviation of at least 3 independent replicates per condition using GraphPad Prism 9. Statistical differences were determined using Student’s unpaired *t*-test for two group comparisons with Welch’s correction. The two-tailed test with P < 0.05 is considered significant in all cases. All statistical analysis was performed in collaboration with the Cornell Statistical Consulting Unit.

## Acknowledgements

We thank all members of the Fischbach lab for valuable discussions of this research; Dr. Siyoung Choi for conducting SEM images; Dr. Lawrence Bonassar for providing materials to produce the bone scaffolds; Dr. Butcher for providing materials and equipment for chick incubation and Shuofei Sun for trainings on handling chicken embryos; Dr. Estroff for her valuable feedback and Stephan Sutter for conducting the Raman analysis; the Cornell Animal Health Diagnostic Core for paraffin embedding, sectioning and H&E staining. Financial support was provided by the National Institutes of Health R01CA259195 and a grant by the Breast Cancer Alliance to C.F; NSF GRFP (DGE-2139899) and Cornell’s Dean’s Excellence and Provost Diversity Fellowship to S.O.K. This work used the Cornell Center for Materials Research (CCMR), which is supported through the NSF MRSEC program (DMR-1719875); and the Cornell University Biotechnology Resource Center (BRC) facilities, including a Zeiss LSM880 confocal/multiphoton microscope (NYSTEM (C029155) and NIH (S10OD018516)), and Bruker Skyscan 1276 micro-CT (NIH S10OD025049).

