## Supplemental for "Directional bone matrix mineralization in the CAM assay is governed by vascular integration and matrix remodeling"

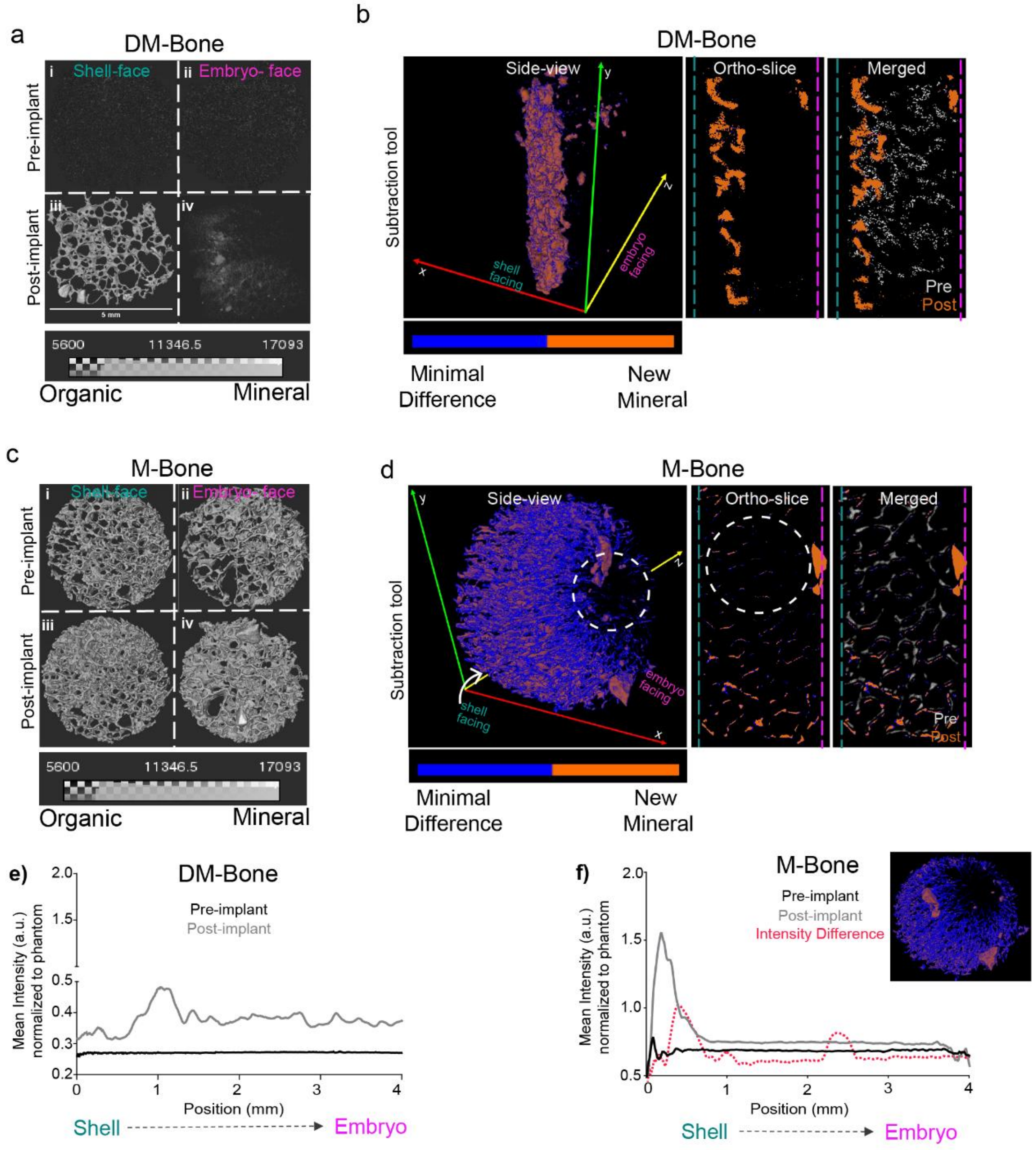


**Supplemental 1. 3D segmentation subtraction tool via Avizo software reveals further matrix remodeling. a,c)** Representative gray-scale images of DM-Bone (a) and M-Bone (c) pre- and post-implant. **b,d)** Image intensity subtraction of DM-Bone (b) and M-Bone (d) pre- from post-implant. Minimal difference in intensity is indicated by blue, greatest difference indicating new mineral deposition is shown in orange. Black regions within M-Bone (d) indicate loss in intensity (white dashed line). Merged scans overlay pre-implant image with post-processing using the subtraction tool. **e,f)** Mean intensity computed per millimeter starting from shell-facing interface to embryo. (f) The intensity difference represents the mean intensity of the subtraction image.

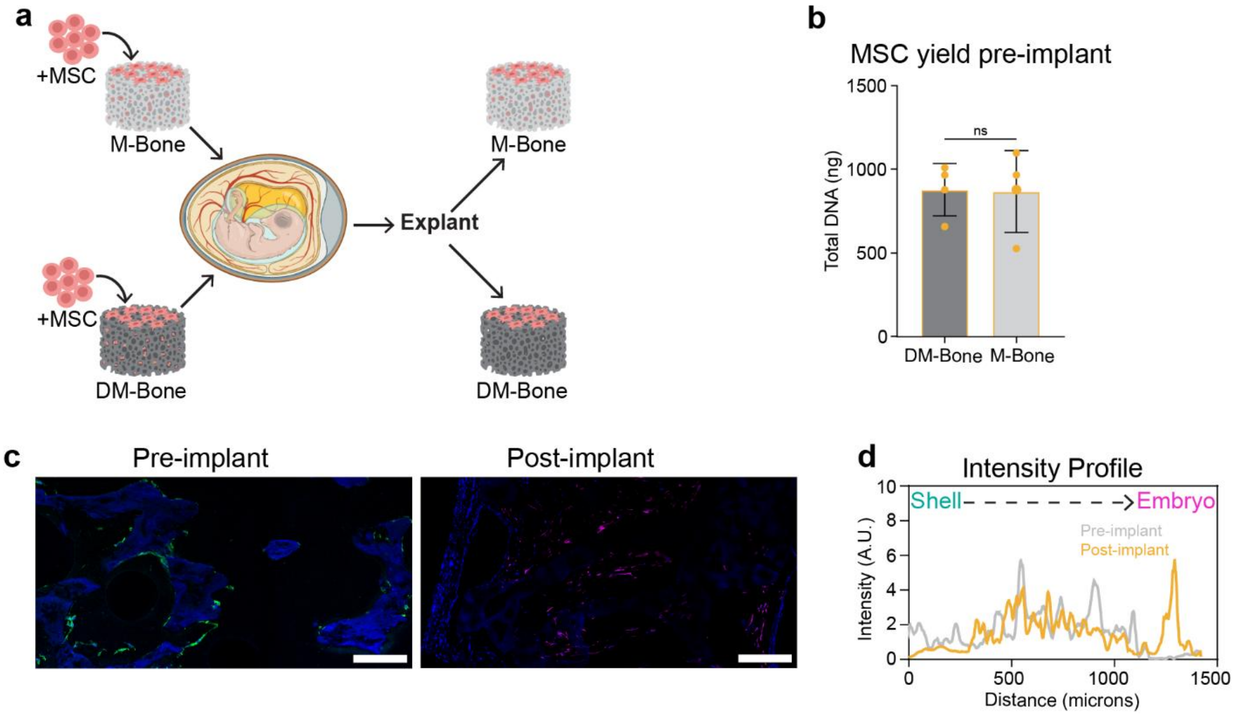


**Supplemental 2. MSCs presence on M-Bone prior and post CAM implantation. a)** Setup procedure for MSC seeding on scaffold systems. **b)** DNA assay confirms similar MSCs yield across both scaffold systems pre-implant. **c)** Fluorescent imaging of MSCs on M-Bone pre- and post-implant (Pre-implant: Phalloidin (green) and DAPI (blue), Post-implant: Human vimentin (magenta) and DAPI (blue)) (scale bar: 200 μm). **d)** Quantification of MSC presence pre- and post-implant was via intensity profile of the phalloidin or vimentin image channel across sample section.

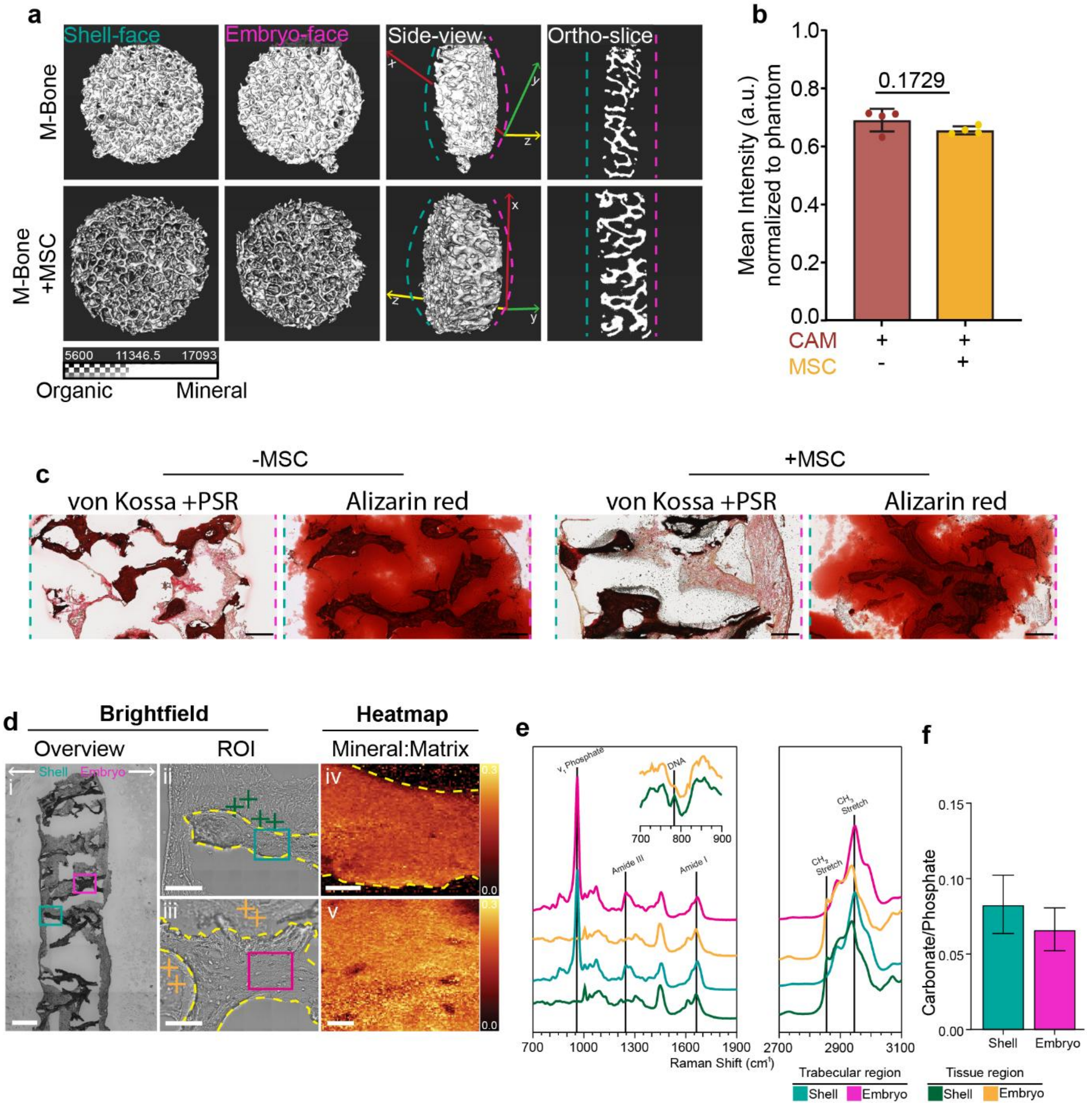


**Supplemental 3. Bone matrix chemical composition remained the same on M-Bone seeded with MSCs**. **a)** Representative µCT images visualizing M-Bone with or without MSCs. **b)** Quantification of mean intensity normalized to phantom of M-Bone with or without MSCs from µCT images. **c)** Von Kossa and PSR or Alizarin Red staining (scale bar: 200 µm). **d)** Raman microscopy: overview brightfield image (i) (scale bar: 500 µm), region of interest (ii) (scale bar: 100 µm), and mineral to matrix ratio heatmap (iii) (scale bar: 20 µm) of M-Bone with MSCs showing high contrast. **e)** Raman spectrum of M-Bone with MSCs. **f)** Quantification of the carbonate and phosphate ratio for shell and embryo region.

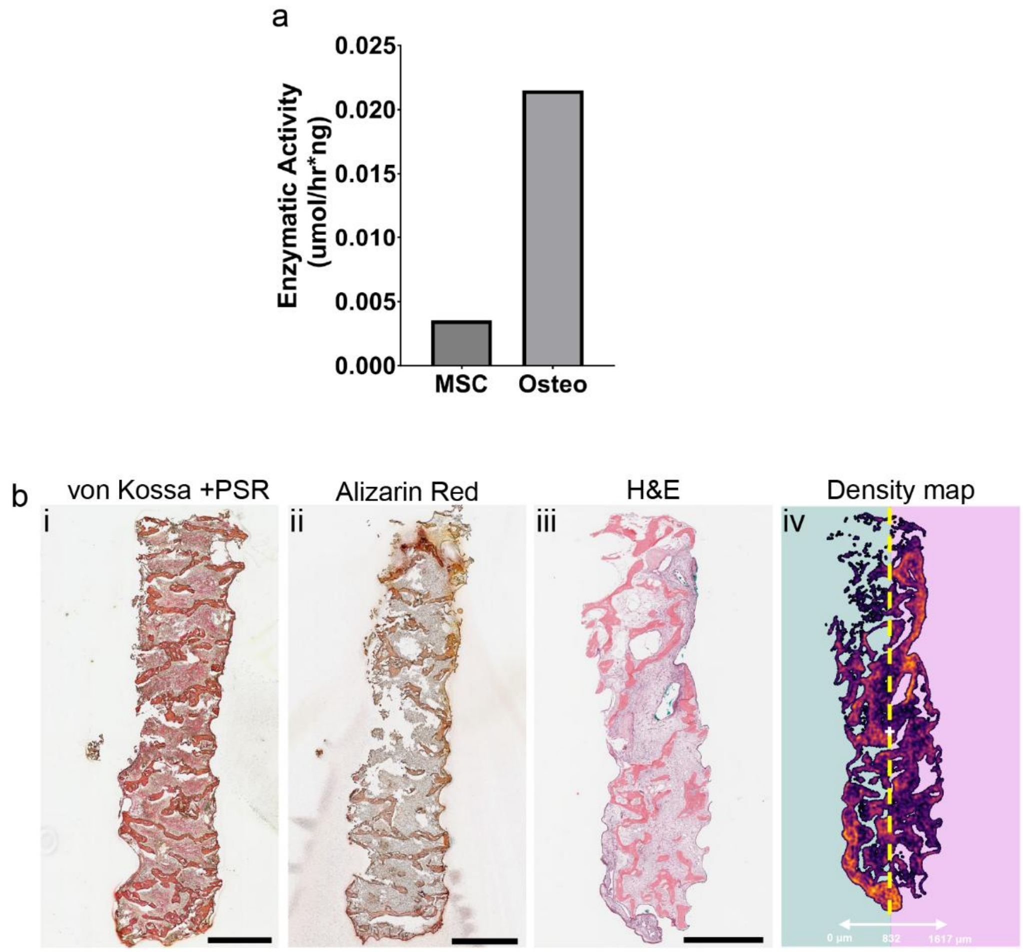


**Supplemental 4. Osteogenic induction of MSCs does not restore mineral deposition. a)** Quantification of ALP activity after 1-week osteogenic induction of MSCs. **b)** 10 μm cryosections of DM-Bone with osteogenically induced MSCs stained with von Kossa and picrosirius red (i), alizarin red (ii), hematoxylin and eosin (iii), and cell density map (iv) (scale bar: 1 mm).

**Supplemental Table 1. Raman band assignments.** Raman shifts and assignments of significant peaks observed in the spectra reported in this work.

| Peak position | Band Assignment | Species |
|---|---|---|
| 776 $cm^{-1}$ | **Nucleic acids**, DNA | Cells |
| 860, 923 $cm^{-1}$ | **Pro**, proline | Collagen |
| 876, 943 $cm^{-1}$ | **Hyp**, hydroxyproline | Collagen |
| 961 $cm^{-1}$ | **$\nu_1(PO_4^{3-})$**, hydroxyapatite | Mineral |
| 1006 $cm^{-1}$ | **Phe**, phenylalanine | Protein |
| 1070 $cm^{-1}$ | **$\nu_1(CO_3^{2-})$**, carbonate ion substitution in HAp | Mineral |
| 1248 $cm^{-1}$ | **Amide III**, collagen triple-helix | Collagen |
| 1340 $cm^{-1}$ | **Amide III**, non-collagenous protein (NCP) | Protein |
| 1615 $cm^{-1}$ | **Nucleic acids**, purines | Cells |
| 1668 $cm^{-1}$ | **Amide I**, protein | Protein |
| 2857 $cm^{-1}$ | **$\nu_{sym.}(CH_2)$**, (lipid) alkyl chains | Cells |

**Supplemental Table 2. Peak integration ranges used in bone parameter ratios.** Integration was done over the listed spectral ranges, using a baseline drawn between the average of either 4 or 0 pixels (flat baseline) to the left and right of the integrated range.

| Raman Peak | Range | L,R Baseline |
|---|---|---|
| **$\nu_1(PO_4^{3-})$**, mineral (phosphate) | 913.5–993.9 $cm^{-1}$ | 4,4 |
| **$\nu_1(CO_3^{2-})$**, carbonate | 1056.2–1091.8 $cm^{-1}$ | 4,4 |
| **$CH_x$ stretch**, organic matrix | 2790.0–3030.0 $cm^{-1}$ | 4,0 |
